# Macrophages discriminate and sort pathogenic bacteria and apoptotic material during dual-target phagocytosis

**DOI:** 10.64898/2026.08.07.743599

**Authors:** Celeste Dea, Paula Arias, Sol L. Moroni, Nicolás N. Rolandelli, Arlinet Kierbel

## Abstract

Macrophages are central to immune homeostasis, clearing both pathogens and apoptotic cells. While these processes share some features, they differ functionally: efferocytosis of apoptotic cells is typically anti-inflammatory, whereas bacterial uptake triggers strong inflammatory responses. During infections, both targets may coexist, yet how macrophages handle such complex particles is poorly understood. Previously, we showed that *Pseudomonas aeruginosa* adheres to apoptotic cells, forming stable composites that can be jointly phagocytosed.

Here, using quantitative confocal and live-cell imaging, we investigate how macrophages engage and internalize these dual targets. We find that macrophages employ distinct strategies depending on target composition. Apoptotic cells alone are engulfed intact within tight, actin-rich membrane cups, whereas bacteria-laden apoptotic cells induce protrusive, actin-driven extensions that navigate along the apoptotic scaffold to access bacteria. Individual bacteria are selectively extracted and internalized, while apoptotic material is internalized in a piecemeal manner. Early phagosome analysis shows that most contain either bacteria or apoptotic material alone, with mixed cargo being less frequent.

These findings demonstrate that macrophages can discriminate and sort the components of complex targets during uptake, revealing a previously unrecognized sophistication in phagocytic processing. This work provides a framework for understanding how innate immune cells integrate pathogen clearance while managing apoptotic material.

## Introduction

Macrophages, as key components of the immune system, play a pivotal role in maintaining homeostasis and responding to various challenges within the body. By virtue of their dynamic and versatile functions, macrophages contribute significantly to the maintenance of tissue integrity and the overall immune response. Their ability to survey the microenvironment is facilitated by surface receptors that constantly monitor for signals indicative of both pathogenic threats and cellular debris. In microbial surveillance, macrophages employ pattern recognition receptors (PRRs) to identify conserved molecular patterns associated with pathogens. This enables them to quickly recognize and respond to invading microorganisms, initiating processes such as phagocytosis to eliminate the threat. But also, macrophages are adept at recognizing apoptotic cells through specific receptors and molecular markers. This process, known as efferocytosis, involves the engulfment and removal of apoptotic cells before they undergo secondary necrosis, preventing the release of potentially harmful cellular contents. Efferocytosis is essential for tissue homeostasis, embryonic development, and immunity. In particular, the uptake and processing of apoptotic cells have a profound influence on the resolution of inflammation.

The particles to be engulfed are covered by different classes of ligands, characteristic of the type of particle. For example, the molecular patterns associated with microbes (Microbe-Associated Molecular Patterns, MAMP), characteristic of certain bacteria or fungi or molecular patterns associated with apoptotic cells (ACAMP) and damage-associated molecular patterns (DAMP) that mark damaged cells but not healthy ones [1] [2] [3] [4]. The receptors present in phagocytes bind to and decode these patterns, triggering specific signaling pathways. While there are aspects of the process that are common, the activation of different receptors leads to particle-specific responses depending on the nature of the particle. For example, the phagocytosis of apoptotic cells and bodies generally triggers an anti-inflammatory response and rapid digestion of particles to prevent the presentation of self-antigens. Contact with bacteria, on the other hand, leads to powerful inflammation and slower degradation, allowing the preservation of peptides for antigen presentation [5] [6] [1]. However, during the infectious process, macrophages may have to interact simultaneously with particles of different origins. In fact, many infections are characterized by an increased number of apoptotic cells. For instance, this is observed in patients with cystic fibrosis (CF), who experience chronic lung infections, particularly with the bacterium *Pseudomonas aeruginosa*. Their airways exhibit an elevated presence of apoptotic cells, attributed in part to dysregulated or excessive apoptosis in CF epithelial cells, along with abnormal efferocytosis [7] [8]. Additionally, environments conducive to *P. aeruginosa* infection, such as wounds and burns, are marked by the presence of a significant number of dead cells, both apoptotic and necrotic [9].

In recent years, numerous examples have emerged that demonstrate an interface between efferocytosis and microbial pathogenesis. In some cases, efferocytosis can serve as a component of a defense mechanism against microbial infections [10] . However, certain pathogens may subvert the efferocytosis process as a strategy during infection [10] [11]. As an illustration of the former scenario, our research has documented that, when infecting polarized epithelial monolayers, *P. aeruginosa* exhibits almost exclusive adherence to extruded apoptotic cells. Notably, extrusion is a mechanism used to eliminate both excess and dying cells from epithelial tissues [12]. The recruitment and adhesion of *P. aeruginosa* to dying extruded cells lead to the formation of bacterial clusters, hereafter referred to as PA–AC clusters, within minutes. Subsequently, neighboring epithelial cells can internalize PA–AC clusters and eliminate the bacteria intracellularly [13] [14]. In a separate study, we observed that macrophages display the ability to ingest PA–AC clusters without compromising their bactericidal efficacy. Additionally, the heightened production of the inflammatory cytokine IL-6 in response to *P. aeruginosa* processing augments the efferocytic capabilities of these phagocytes [15]. Hence, the internalization and digestion of pathogenic bacteria and apoptotic cells by macrophages should not automatically be regarded as distinct and independent processes. This underscores the intricate nature of interactions between the immune system and the presence of pathogens. Within this context, numerous questions arise: Does the presence of two different targets modify the mechanism of uptake by macrophages? Are macrophages able to distinguish the different targets? How are dual targets processed intracellularly?

In the current study, we exposed mouse bone marrow-derived macrophages as well as human derived macrophages to PA–AC clusters, and elucidated how the same phagocytic cell manages the uptake and processing of two distinct types of particles simultaneously.

## Results

### Differential engulfment of apoptotic cells and *P. aeruginosa* by BMDMs

We investigated the modes of engulfment used by bone marrow–derived macrophages (BMDMs) during interactions with *P. aeruginosa* clusters on apoptotic cells (PA–AC clusters). As a first step, we examined the phagocytic morphology when each target was presented individually.

Following incubation with UV-induced apoptotic cells, BMDMs were examined by confocal microscopy and were observed to exhibit a predominantly rounded morphology. Apoptotic cells became closely associated with the BMDM cell body, fully encased within a tight, actin-rich membrane cup (Fig 1A), as previously described [16].

**Fig 1.**
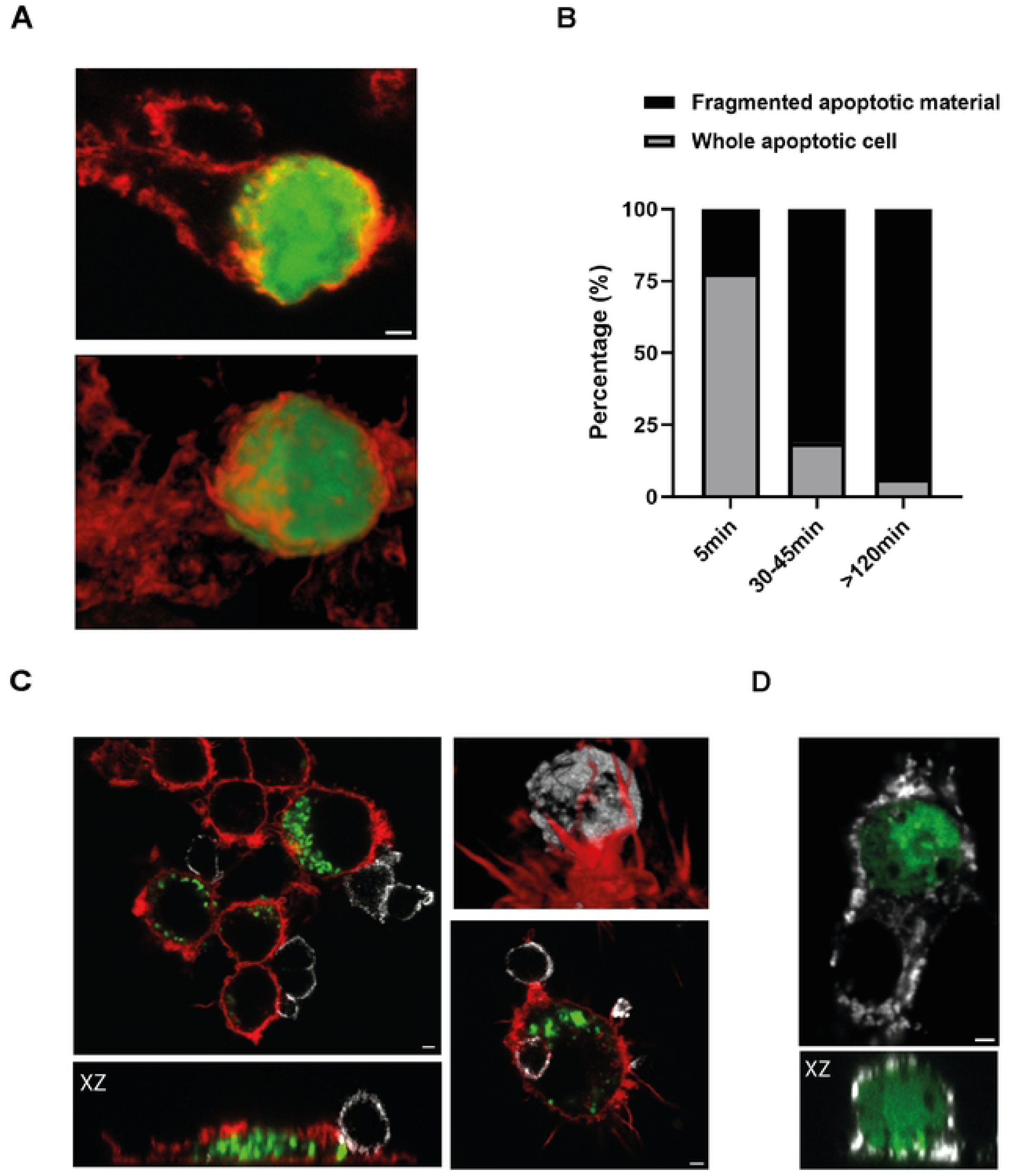
BMDMs capture intact apoptotic cells and process them into intracellular fragments. BMDMs were coincubated with apoptotic cells labeled with CFSE (green) or Annexin V–Alexa Fluor 647 (gray) for the indicated times and analyzed by confocal microscopy A) An apoptotic cell undergoing internalization is fully enclosed within a tight, actin-rich membrane cup. The lower panel shows a 3D reconstruction of the upper panel image. B) BMDMs were exposed to apoptotic cells at a 1:3 apoptotic cell-to-macrophage ratio for varying times, and the percentage of macrophages containing intact or fragmented apoptotic material was quantified. C) BMDMs underwent two consecutive rounds of efferocytosis with apoptotic cells labeled with different fluorescent markers. Apoptotic cells were exposed for 45 min in the first round (green), washed, and 120 min later a second round (gray) was performed; samples were fixed 5 min afterward. The upper right panel shows a 3D reconstruction of part of the image in the lower right panel. D) BMDMs were exposed to CFSE-labeled apoptotic cells for 5 min and stained for LAMP1 (gray). Red: F-actin. XZ: longitudinal reconstruction. Scale bars: 2 μm.

Studies investigating phosphatidylserine-mediated phagocytosis or efferocytosis have revealed diverse mechanisms. Both *in vivo* and *in vitro* studies have reported piecemeal uptake, a process in which portions of an apoptotic cell are ‘nibbled off’ and internalized [17] [18] [19]. Conversely, as previously mentioned, other studies have described the engulfment of intact apoptotic cells [20]. While our preceding observations indicate that apoptotic cells can be internalized intact, BMDMs exposed to apoptotic cells for longer times displayed apoptotic material compartmentalized within multiple vesicles. This raises the possibility that, in addition to whole-cell engulfment, piecemeal uptake may also occur in this system. Alternatively, this phenomenon may reflect the internalization of intact apoptotic cells followed by subsequent intracellular fractioning [21]. To evaluate this, we exposed BMDMs to apoptotic cells at a ratio of one apoptotic cell per three macrophages for varying durations and quantified the percentage of macrophages containing either a whole internalized apoptotic cell or fragmented apoptotic material. We found that shorter exposure times yielded a higher proportion of macrophages containing intact apoptotic cells, whereas longer exposures increased the proportion containing fragmented apoptotic material. (Fig 1B). Fig 1C shows confocal micrographs of BMDMs subjected to two consecutive rounds of efferocytosis, each using apoptotic cells labeled with a different fluorescent marker. In the first round, BMDMs were exposed to CFSE-labeled apoptotic cells for 45 minutes; cells were then washed, and 120 minutes later a second round was initiated using Annexin V–Alexa Fluor 647–labeled apoptotic cells. Samples were fixed 5 minutes after the second incubation. CFSE-labeled apoptotic cells from the first round appear as intracellular fragments. In contrast, 5 minutes into the second incubation, Annexin V–labeled apoptotic cells are found either at the BMDM surface, often surrounded by filopodia (Fig 1C, upper-right panel), or already internalized while still morphologically intact (Fig 1C, bottom-right panel). Thus, our results are consistent with the possibility that apoptotic cells may be internalized intact and subsequently fragmented intracellularly.

By five minutes after entry into the cell, apoptotic cells were already enclosed within lysosomal-associated membrane protein 1 (LAMP1)-positive vesicles (Fig 1D) [22].

We then exposed BMDMs to *P. aeruginosa* for short times. BMDMs were observed more extended, forming elongated membrane protrusions. Bacteria were often observed within these protrusions (Figs 2A, B and D). Live microscopy revealed continuous membrane ruffling along these extensions, a well-established mechanism that facilitates bacterial capture and internalization (S1 Video). Protrusions can be very long, with an average length of 34.1 ± 13.1 microns, and they align with microtubules, as revealed by staining with an α-tubulin antibody (Fig 2B). Macrophages were stained with an antibody against LAMP1. In both uninfected and infected cells (Figs 2B and C respectively), LAMP1-positive vesicles were observed not only in the perinuclear region but also toward the cell periphery, aligned along the microtubular network, consistent with previous reports [23] [24]. Multiple LAMP1-positive vesicles were also present within membrane protrusions, positioned along microtubule fibers. Following phagocytosis, bacteria were rapidly detected within LAMP1-positive vesicles (Fig 2D). Although we cannot directly assess their dynamics, the peripheral and protrusion-localized positioning of these vesicles suggest that macrophages may maintain a pool of LAMP1-positive compartments poised for rapid fusion with phagocytic cargo.

**Fig 2.**
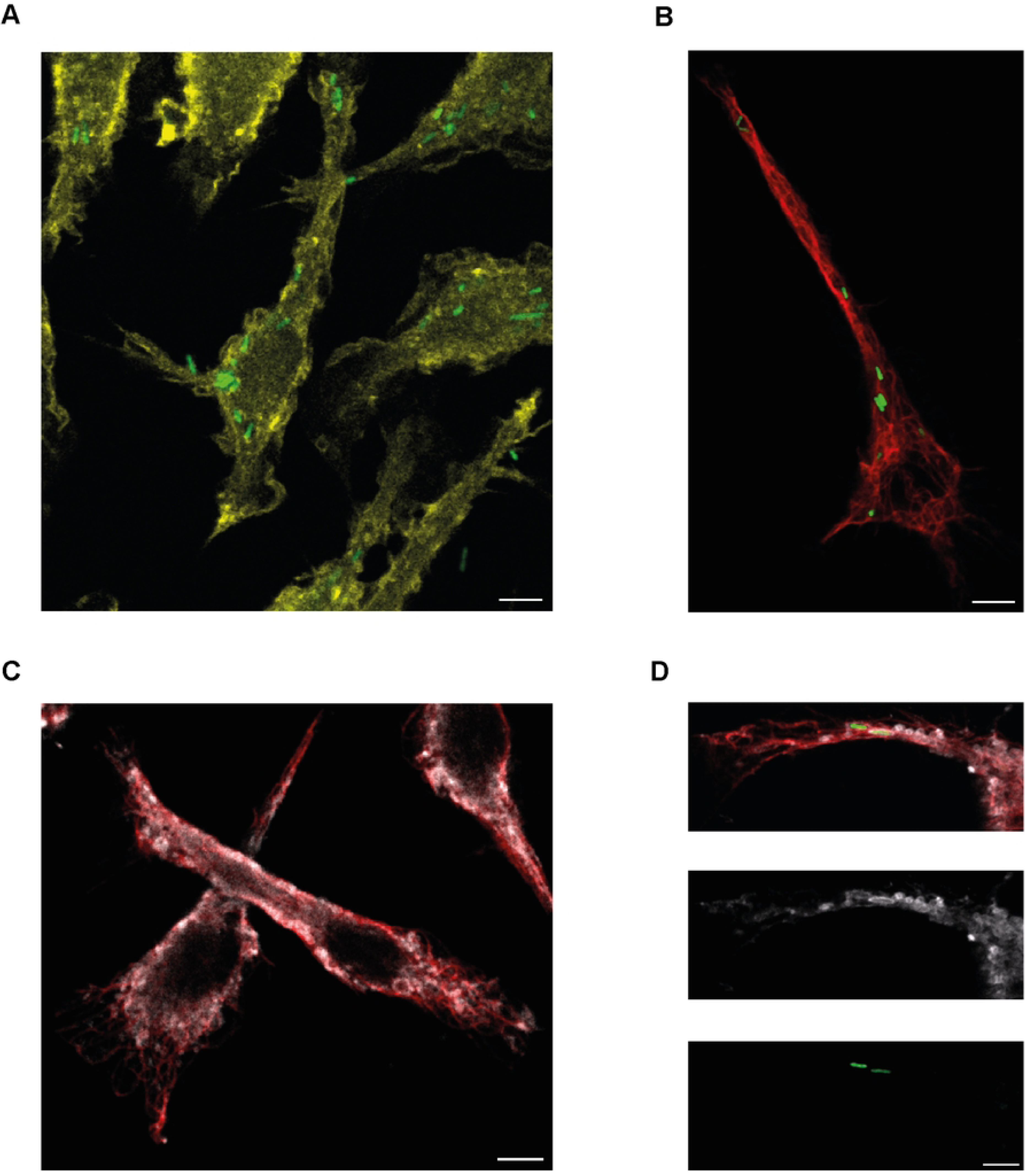
Engulfment of *P. aeruginosa* and localization to LAMP1-positive, microtubule-associated vesicles. BMDMs were co-incubated with GFP–*P. aeruginosa* (green) for 15 minutes and examined by confocal microscopy (A, B, D). Panel C shows non-infected BMDMs. Yellow: F-actin, red: α-tubulin; gray: LAMP1. Scale bars: 5 μm.

In summary, BMDMs predominantly internalize apoptotic cells as intact structures within tight, actin-rich membrane cups, followed by intracellular fragmentation. Apoptotic cell containing-phagosomes are LAMP1-positive at early time points. *P. aeruginosa* can be engulfed by macrophages membrane extensions and is also detected early in LAMP1-positive vesicles associated with microtubules, suggesting trafficking along microtubule-dependent pathways.

### Stable association of *P. aeruginosa* with the apoptotic surface in PA–AC clusters

In our previous work, we characterized the dynamics of PA–AC cluster formation with apoptotic cells extruded from epithelial monolayers and found that bacterial attachment is initially reversible. In this early phase (∼15 min), *P. aeruginosa* primarily attaches to the surface of apoptotic cells via the bacterial pole [14]. In the present study, PA–AC clusters were generated *in vitro* as previously described [15]. Bacteria were incubated with UV-induced apoptotic J774 cells for 1 hour. Within these PA–AC clusters, *P. aeruginosa* is no longer pole-attached; instead, the bacteria become embedded within the apoptotic material and align along their longitudinal axis, indicating a transition from transient surface attachment to stable incorporation within the cluster (Fig 3). To assess the stability of PA–AC clusters, we tracked the position of individual bacteria attached to the surface of apoptotic cells. A total of 115 bacteria from 10 independent PA– AC clusters were tracked. Of these, 100 bacteria remained associated with the apoptotic cell throughout the entire observation period, corresponding to 87% of the tracked bacteria. These findings indicate substantial stability of PA–AC clusters following the initial incubation period, with most bacteria remaining embedded within apoptotic cells over time (Fig 3B and S2 Video).

**Fig 3.**
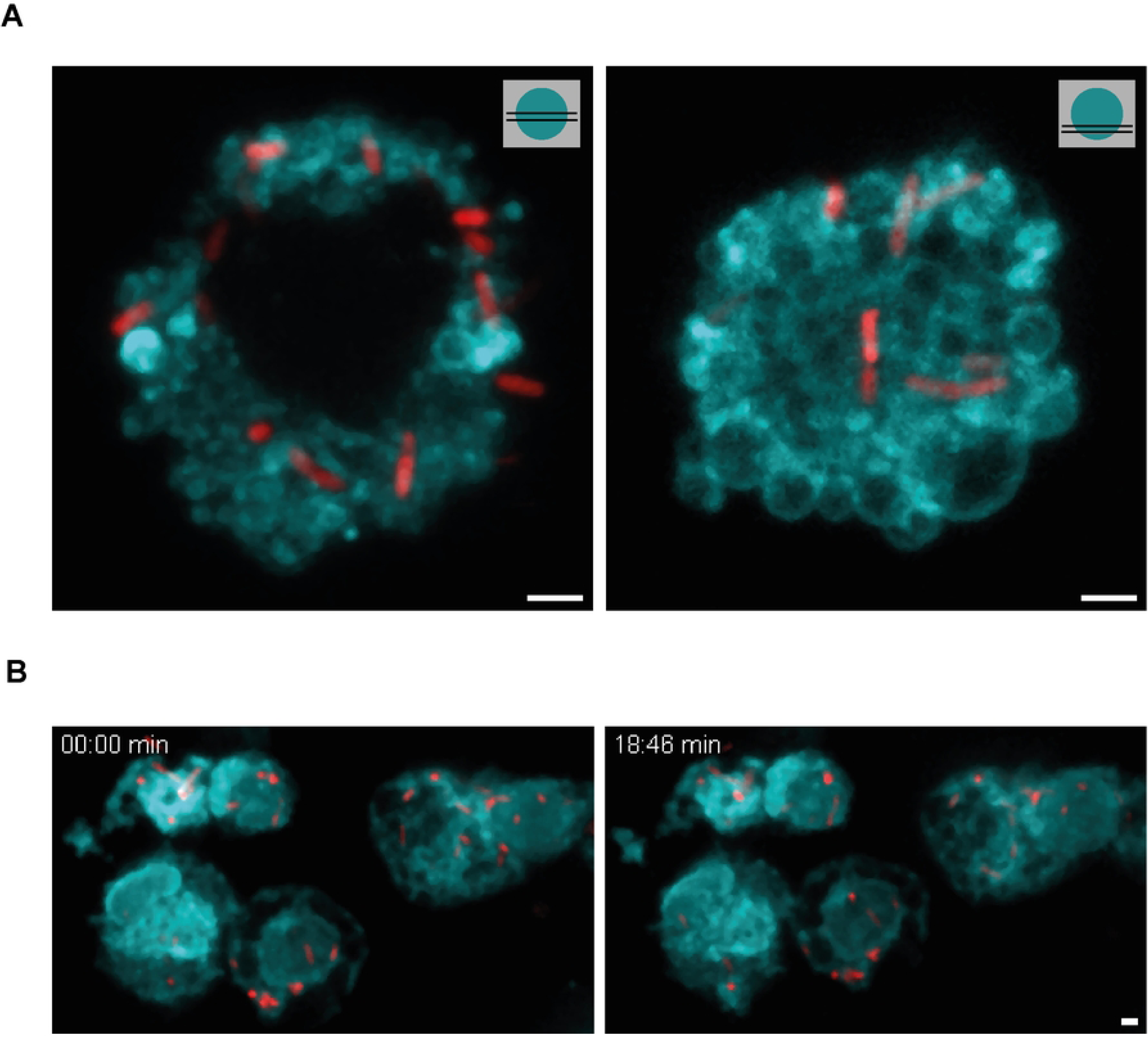
Stability of PA–AC clusters. A) Maximum-intensity projections of PA–AC clusters at the equatorial (left) and polar (right) regions of the apoptotic cell. Cyan: AnnexinV-Alexa 647-labeled apoptotic cells; red: GFP-*P. aeruginosa* B) Snapshots (maximum-intensity projections) from z-stack time-lapse imaging (∼20 min), showing that bacteria maintain their positions within PA–AC clusters. Cyan: Cell Trace-labeled apoptotic cells, red: mCherry-*P. aeruginosa*. Scale bars: 2 μm.

### Shift in apoptotic cell uptake mode in the presence of *P. aeruginosa*

In our previous studies, we demonstrated that, when presented with PA–AC clusters, BMDMs are capable of internalizing both components, *P. aeruginosa* and apoptotic cells [15]. We are now interested in investigating how macrophages manage the uptake of this dual target. We exposed both BMDMs and human monocyte-derived macrophages to PA–AC clusters, for shorter durations. Fig 4 illustrates a notable distinction in macrophage behavior when internalizing PA– AC clusters compared to apoptotic cells alone. While apoptotic cells alone, as shown, can be internalized intact, bacteria-laden apoptotic cells undergo fragmentation during internalization by the macrophages. In all observed instances of partial PA–AC cluster internalization (BMDMs or human-derived macrophages incubated with PA–AC clusters for 15 minutes), apoptotic material was consistently found fragmented within LAMP1-positive vesicles (Fig 4A, S1 Fig A, and S3 Video). This process more closely resembles a piecemeal mode of uptake and suggests a shift in the internalization mechanism triggered by the presence of bacteria. To confirm that these fragmentation events occur before complete internalization of PA–AC clusters, we differentially labeled extracellular material after 30 min of incubation. Annexin V stained only the non-fragmented apoptotic material remaining outside the macrophage, whereas intracellular CFSE-positive fragments were not labeled (Fig 4B and S4 Video). Fragmentation was observed in all partially engulfed apoptotic cells analyzed across three independent experiments, indicating that apoptotic cell fragmentation consistently occurs before complete engulfment. At 30 min, BMDM interacting with partially engulfed apoptotic cells contained an average of 8.4 ± 5.9 intracellular CFSE-positive fragments (mean ± SD). Similarly, anti-*Pseudomonas* staining labeled extracellular bacteria associated with the cluster but not bacteria contained within intracellular compartments (Fig 4C and S5 Video). Together, these findings confirm that PA–AC cluster uptake occurs progressively, with extracellular cluster material coexisting with intracellular fragmented cargo during the same internalization event.

**Fig 4.**
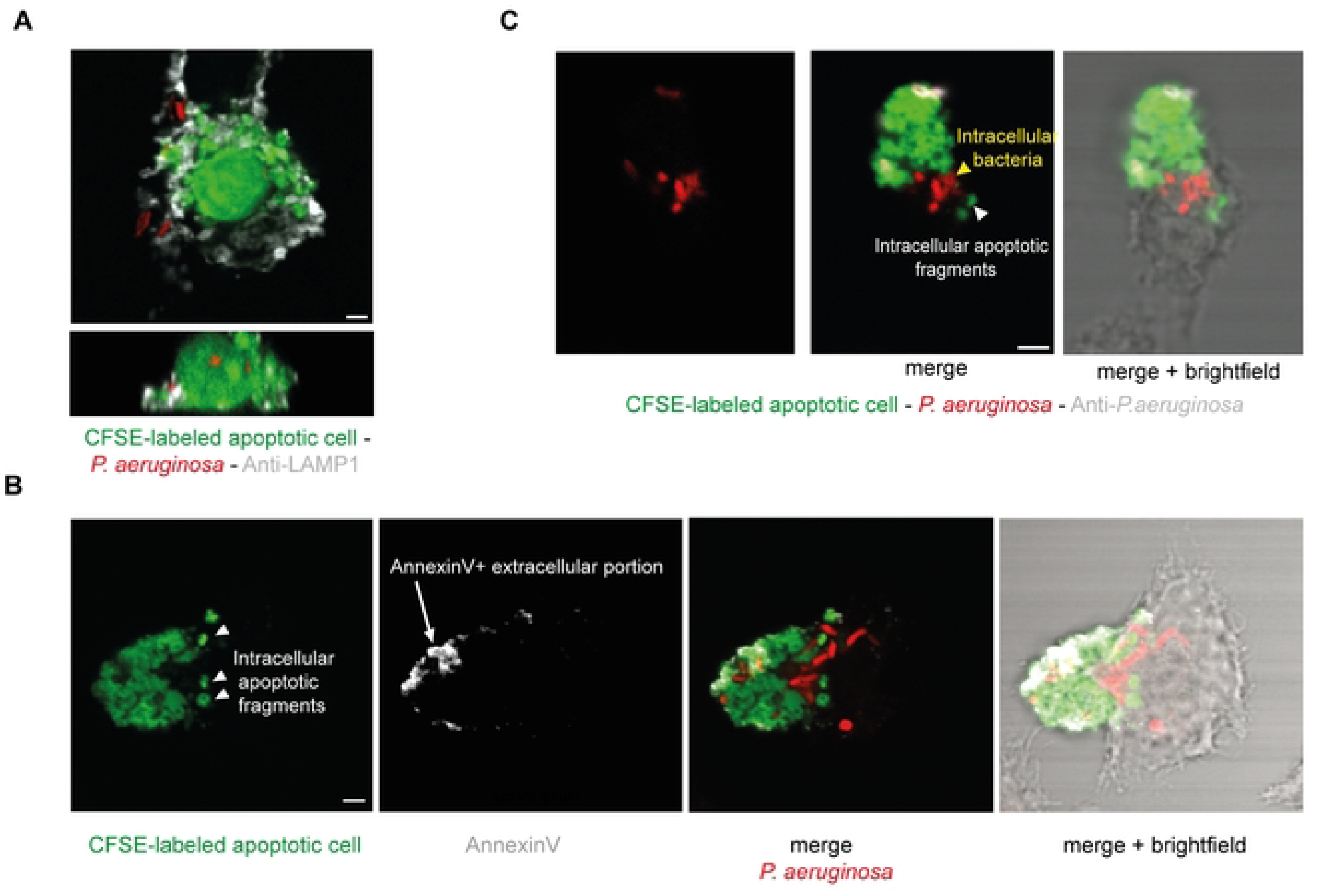
Bacteria Shift Apoptotic Cell Uptake Mode. BMDMs were exposed to PA–AC clusters for the indicated times, treated as described, and analyzed by confocal microscopy. **(A–C)** Representative images of partially internalized PA– AC clusters. In all cases, apoptotic cell fragmentation is evident while part of the PA–AC cluster remains extracellular. **(A)** Samples were fixed, permeabilized, and labeled with an anti-LAMP1 antibody. The upper panel shows a maximum-intensity projection, and the lower panel an orthogonal view of a partially internalized PA–AC cluster after 15 min of coincubation. Gray, LAMP1. **(B)** Before fixation, samples were labeled with Annexin V–Alexa Fluor 647. A representative confocal optical section is shown. Annexin V labeled only the non-fragmented apoptotic material that remained extracellularly accessible, whereas intracellular CFSE-positive fragments were not labeled. Coincubation time: 30 min. **(C)** Non-permeabilized samples were labeled with an anti-*Pseudomonas aeruginosa* antibody (Gray). A representative confocal optical section is shown. The antibody labeled extracellular bacteria associated with the apoptotic cell, but not bacteria that had already been internalized. Coincubation time: 30 min. Scale bars 2 μm.

We then asked whether the presence of bacteria—capable of altering the mode of apoptotic cell uptake—would affect the efferocytic outcome compared with apoptotic cells alone. In our experimental setup, BMDMs were exposed either to apoptotic cells alone or to an equivalent number of apoptotic cells carrying bacteria for 30 min, followed by fixation. Subsequent confocal microscopy and image-based quantification of internalized apoptotic material, as described in the Methods section, showed that BMDMs internalized less apoptotic material in the presence of bacteria (S1 Fig B). The shift in the apoptotic cell uptake mode observed in the presence of bacteria may contribute to a reduction in overall uptake efficiency. It is noteworthy that our previous findings showed no significant change in the efficiency of phagocytosis and/or killing of *P. aeruginosa* in the presence of apoptotic cells [15], whereas the present results show reduced internalization of apoptotic material under the same conditions.

### PA–AC cluster uptake depends on actin polymerization and microtubule integrity

Closer examination of actin distribution reveals distinct differences: unlike the case with apoptotic cells alone, macrophages interacting with PA–AC clusters fail to form actin-rich membrane cups that tightly enclose the entire target. Instead, macrophages appear to adopt membrane structures resembling those observed during the uptake of bacteria alone, with elongated actin-rich protrusions that either extend around (Fig 5A, lower panel and S6 Video) or extend through existing gaps in the apoptotic cell (Fig 5A, upper panel and S7 Video), enabling access to the bacteria within the clusters. The extensions often displayed features indicative of membrane ruffling. We quantified the presence and number of the extensions during the early stages of macrophage–PA–AC cluster interactions and assessed whether their tips contacted or enclosed bacteria. Approximately 75% of the extensions examined met this criterion, both in human macrophages and BMDMs. We also observed extensions that enclosed apoptotic material, suggesting that ruffling activity triggered by bacterial presence can capture small fragments of apoptotic cells. S2 Fig shows the distribution of actin-rich protrusions per macrophage, indicating that the majority of cells had protrusions, often more than one per cell.

**Fig 5.**
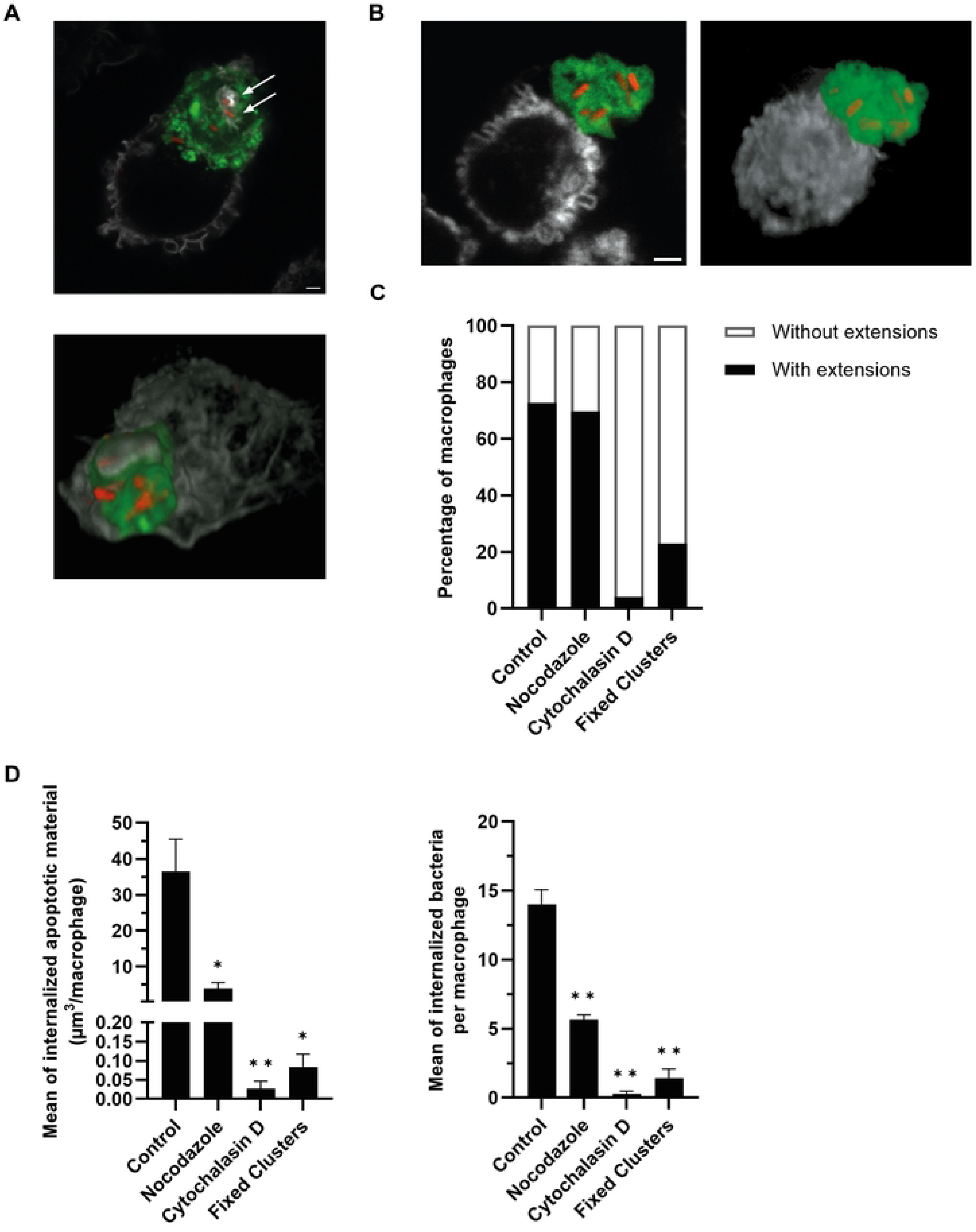
Protrusion-Mediated PA–AC Cluster Uptake Depends on Cytoskeletal Integrity. BMDMs or human monocyte-derived macrophages were exposed to PA–AC clusters under the indicated conditions and analyzed by confocal microscopy. In representative micrographs, F-actin is shown in gray, CFSE-labeled apoptotic material in green, and mCherry-*P. aeruginosa* in red. **A)** A human macrophage (upper panel) and a BMDM (lower panel; 3D reconstruction) were exposed to PA–AC clusters for 15 or 5 min, respectively. Actin-rich elongated protrusions, either projecting through pre-existing gaps in the apoptotic cell (upper panel, white arrows) or extending around the apoptotic cell (lower panel), provide access to bacteria within the clusters. **B)** BMDMs were treated with 2 μM cytochalasin D and exposed to PA–AC clusters for 15 min. Cytochalasin D treatment abolished the formation of actin-rich protrusions. The right panel shows a 3D reconstruction of the image on the left. **C)** Percentage of BMDMs exhibiting actin-rich protrusions after 15 min of incubation with PA–AC clusters under control conditions or following treatment with 2 μM cytochalasin D, 5 μM nocodazole, or exposure to fixed PA–AC clusters. Cytochalasin D and nocodazole were added 1 h before PA–AC cluster exposure and maintained throughout incubation. **D)** Quantification of internalized apoptotic material (left panel) and bacteria (right panel) per macrophage after the treatments described in (C). Statistical analysis was performed using multiple unpaired t-tests comparing each treatment with the control. Data represent macrophages analyzed from three independent experiments and are shown as mean ± SEM. * indicates *p* < 0.05 and ** indicates *p* < 0.01. Scale bars: 2 μm.

To determine whether actin polymerization is required for the formation of these protrusions and for PA–AC cluster uptake, BMDMs were treated with cytochalasin D before and throughout incubation with PA–AC clusters. Inhibition of actin polymerization nearly abolished the formation of actin-rich protrusions and the internalization of both bacteria and apoptotic material, as assessed at 15 and 30 min, respectively (Fig 5B-D). As protrusions were found to align with microtubules (Figs 2B–D), we next examined the effect of microtubule depolymerization. BMDMs were treated with nocodazole before and throughout incubation with PA–AC clusters. Although nocodazole treatment did not affect the proportion of macrophages exhibiting actin-rich protrusions (Fig 5C), it reduced the internalization of both apoptotic material and bacteria (Fig 5D), indicating that microtubules contribute to efficient PA–AC cluster internalization.

Together, these results show that actin polymerization is essential for protrusion formation and PA–AC cluster internalization, while microtubules contribute to efficient uptake. Further studies will be required to determine the mechanisms regulating actin remodeling and the functional contribution of microtubules to PA– AC cluster internalization.

To further assess whether features associated with live bacteria contribute to the uptake process, we exposed macrophages to fixed PA–AC clusters. Compared with unfixed PA–AC clusters containing live bacteria, fixed clusters induced fewer actin-rich protrusions and strongly reduced the internalization of both apoptotic material and bacteria. These results suggest that features associated with live bacteria are required for efficient protrusion formation and PA–AC cluster internalization (Figs 5C and D).

### Macrophages sort bacterial and apoptotic material during uptake

As described, during the early stages of contact with PA–AC clusters, the majority of macrophage membrane extensions had their distal ends in contact with bacteria or already contained bacteria within them. To estimate the relative availability of apoptotic material and bacteria within the *in vitro*-generated PA–AC clusters used in this study, we quantified the surface area of each component. The estimated surface area of apoptotic material was 29.0 ± 13.8-fold greater than the estimated bacterial surface area (mean ± SD, *n* = 17 PA–AC clusters). Thus, despite the overwhelming predominance of apoptotic material within the clusters, macrophage membrane extensions were preferentially associated with bacteria during the initial stages of cluster engagement, suggesting that macrophages preferentially recognize bacteria rather than interacting solely according to target surface availability.

To better understand this process, we followed it in real time using the RAW 264.7 macrophage cell line, transiently transfected with plasma membrane marker PM-GFP [25]. Time-lapse imaging reveals that membrane extensions from the macrophage spread over the surface of the PA–AC cluster. Upon contact with individual bacteria on the cluster surface, ruffling activity intensifies, and the membrane surrounds the bacteria and retracts, pulling them out of the cluster and resulting in their internalization inside the macrophage (S8 Video) As shown earlier, *P. aeruginosa* attaches stably to the apoptotic surface. This suggests that the extending macrophage membrane exerts traction forces to detach the bacteria. To visualize the actin dynamics associated with bacterial extraction and subsequent internalization, RAW 264.7 macrophages stably expressing LifeAct-RFP were imaged by live-cell microscopy, revealing extensive actin remodeling during these events. An actin-rich protrusion extends from the macrophage through openings within the PA–AC cluster and reaches a bacterium within the cluster. A phagocytic cup then forms around the bacterium, followed by intense actin-rich membrane ruffling similar to that observed in S8 Video. These coordinated actin-dependent remodeling events contribute to bacterial extraction from the cluster and subsequent internalization by the macrophage (S9 Video). To assess whether the described uptake process leads to sorting of material inside macrophages, BMDMs or human-derived macrophages were co-incubated with PA–AC clusters for 15 or 30 min, after which samples were fixed and stained for LAMP1. A PA–AC-cluster/macrophage ratio of 1:3 was used to ensure that each macrophage interacted with no more than one cluster and to minimize the presence of unattached, free-swimming bacteria. Confocal microscopy Z-stacks were analyzed, and the number of vesicles containing bacteria alone, apoptotic material alone, or mixed cargo (both bacteria and apoptotic material within the same vesicle) was quantified. We found that the majority of vesicles (approximately 80%) contained a single cargo type, whereas only approximately 15–20% contained mixed cargo. Strikingly, this distribution was very similar in BMDMs and human macrophages at both 15 and 30 min. (Fig 6).

**Fig 6.**
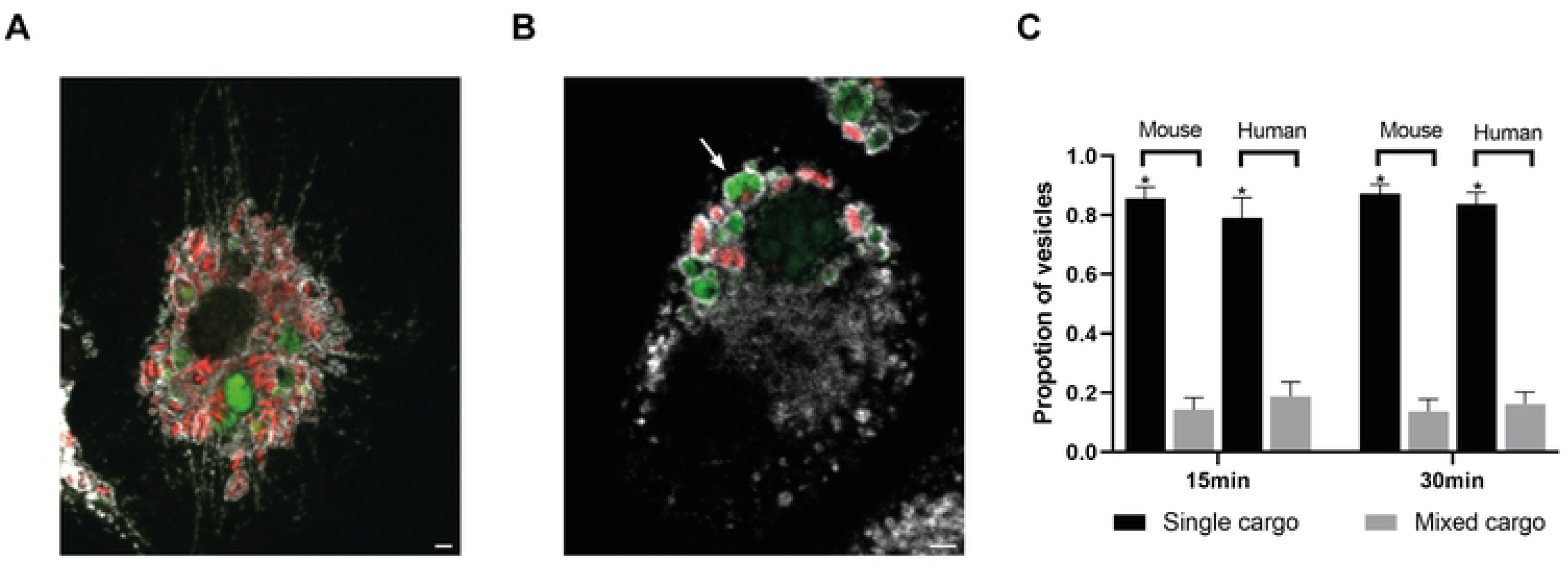
Macrophages Sort the Content of PA–AC Clusters During Uptake. BMDMs or human-derived macrophages were co incubated with PA–AC clusters for 15 or 30 minutes. Confocal Z stacks were analyzed to quantify vesicles with distinct cargo identities (bacteria alone, apoptotic material alone, or mixed cargo). A) Representative image of a BMDM at 30 min (maximum intensity Z projection). B) Representative image of a human macrophage at 30 min; white arrow points to mixed cargo vesicle. C) Quantification of vesicle types per macrophage. Statistical analysis was performed using multiple paired t-tests comparing mixed- and single-cargo conditions within each species and time point. Data are shown as mean ± SEM from three independent experiments. * indicates *p* < 0.001.

Taken together, these results indicate that macrophages efficiently sort the cargo components and support the initiation of cargo segregation during uptake. However, they do not exclude the possibility that subsequent intracellular processing also contributes to the final distribution of cargo. Thus, macrophages can discriminate between distinct components of complex targets.

## Materials and methods

### Reagents and plasmids

The following reagents were used: anti-LAMP1 antibody (clone 1D4B; DSHB); Anti-LAMP1 H4A3 DSHB (Hybridoma Bank); α-tubulin YOL1/34 (Santa Cruz), anti-*Pseudomonas* (Abcam, ab68538), Alexa Fluor 647 goat anti-rat IgG and CFSE (Invitrogen); Nocodazol (Sigma-Aldrich), Alexa Fluor 647–conjugated phalloidin and Cytochalasin D (Life Technologies); Dulbecco’s Modified Eagle Medium (DMEM) and RPMI 1640 (Gibco); recombinant murine M-CSF (PeproTech); and PE-conjugated anti-mouse CD11b (BioLegend). PM-GFP encodes the myristoylation/palmitoylation sequence from Lyn fused to GFP in pcDNA3 vector. PM-GFP was a gift from Tobias Meyer (Addgene plasmid # 21213; http://n2t.net/addgene:21213 ; RRID:Addgene_21213) [25].

### Cell culture

Murine macrophage cell lines J774A.1 and RAW264.7, as well as bone marrow– derived macrophages (BMDMs) from BALB/c mice, were cultured at 37 °C in a humidified atmosphere with 5% CO₂. J774A.1 cells and BMDMs were maintained in RPMI 1640 supplemented with 10% fetal bovine serum (FBS), 1% antibiotic– antimycotic, 2 mM sodium pyruvate, and 2 mM L-glutamine. RAW264.7 cells and RAW cells stably expressing LifeAct-RFP (Dr. Rene Harrison, University of Toronto) were cultured in DMEM supplemented with 10% FBS. All animal procedures were conducted in accordance with institutional guidelines and approved by the Ethics Committee for the Care and Use of Experimental Animals of the Universidad Nacional de San Martín (Permit number 20/2024, CICUAE-UNSAM). Mice were euthanized by CO₂ inhalation followed by cervical dislocation. Primary Human macrophages were cultured in RPMI-1640 supplemented with 10% HI-FBS, 100 U mL−1 penicillin, 100μg mL−1 streptomycin, 25 ng mL−1 <u>h</u>uman <u>M</u>acrophage <u>C</u>olony <u>S</u>timulating <u>F</u>actor (hM-CSF; Invitrogen). All procedures were approved by the University of Toronto Health Sciences Research Ethics Board, Human Protocol #: 00043264. Written informed consent was obtained from all participants prior to blood sample collection, with recruitment conducted between 31 January and 31 July 2025.

### BMDM and human macrophages preparation

Bone marrow–derived macrophages (BMDMs) were generated from fresh bone marrow cells using standard procedures [26]. Cells were resuspended in RPMI complete medium supplemented with 20 ng/mL M-CSF and seeded in 100-mm culture dishes. On days 3 and 6, half of the medium was replaced with fresh supplemented RPMI. On day 8, macrophages were detached by gentle scraping with ice-cold PBS. Flow cytometric analysis confirmed that >98% of cells were CD11b-positive. To obtain relatively non-polarized primary human macrophages, human monocytes were isolated from the blood of healthy donors by density-gradient separation with LymphoprepTM (STEMCELL Technologies), after which human monocytes were then separated by adherence and incubated in RPMI complete medium with 25 ng hM-CSF for 7-10 days at 37 °C in a CO2 incubator. Culture media was replaced with fresh hM-CSF containing media every other day.

### Generation of apoptotic cells

Apoptotic cells (ACs) were generated as described previously [15]. Briefly, confluent J774 cells were resuspended in serum-free RPMI, optionally labeled with 2.5 μM CFSE, and seeded in 100-mm plates. Cells were irradiated with short-wave UV (50 mJ/cm², 1 min) using a UVP CL-100 Crosslinker, then incubated 16 h in serum-free medium. Non-adherent ACs were collected, and apoptosis was confirmed by flow cytometry.

### Bacterial culture and PA–AC cluster generation

*Pseudomonas aeruginosa* strain K (PAK) was grown overnight in LB medium at 37 °C with shaking. Stationary-phase bacteria were used for experiments. For fluorescence microscopy, PAK carrying either the mCherry plasmid (pMP7605-mCherry, kindly provided by Dr. Lagendijk[27] or a GFP plasmid (pBBR1MCS-5 + gfpmut3[14]) was used. For infections with bacteria alone, the multiplicity of infection (MOI) was 40. To generate PA–AC clusters, bacteria and apoptotic cells were co-incubated at a ratio of 15:1 bacteria:AC for 1 h at 37 °C in RPMI medium prior to addition to macrophages. Control apoptotic cells were incubated under identical conditions and for the same duration, without bacteria.

For experiments using fixed PA–AC clusters, clusters were fixed with 4% paraformaldehyde for 20 min at room temperature. Residual aldehydes were subsequently quenched with 15 mM NH₄Cl for 15 min.

### Microscopy studies

BMDMs or human macrophages were cultured on glass coverslips in 24-well plates at a density of 2 × 10⁵ cells per well. Following the indicated treatments, samples were fixed with 4% paraformaldehyde in PBS for 15 minutes at room temperature, in the dark. BMDMs were blocked with 2% BSA in PBS and permeabilized with 0.3% Triton X-100. When necessary, samples were incubated overnight at 4°C with the primary antibody, followed by a 1-hour incubation at 37°C with the secondary antibody and phalloidin. For immunostaining, human macrophages were permeabilized with cold methanol. For LAMP-1 staining, samples were blocked with 5% skim milk in PBS prior to incubation with the primary antibody. For F-actin staining with phalloidin, cells were permeabilized with Triton X-100 and blocked with 5% skim milk before staining.

For immunofluorescence analysis, images were acquired using either a Stellaris 5 laser-scanning confocal microscope (Leica Microsystems) mounted on a Leica DMi8 inverted platform and controlled with Leica Application Suite X (LAS X) software, equipped with a 63× oil-immersion objective (NA = 1.4), or an Olympus FV1000 confocal laser-scanning microscope with a PlanApo N 60X oil objective (1.42 NA). Images were acquired in the XY plane along the Z-axis using a z-step increment of 0.22 μm (Z-stack). Five to seven fields were randomly selected from each coverslip, and all cells within a given field were included in the image analysis. Live-cell imaging of RAW macrophages was performed using a WaveFX-X1 spinning-disk confocal system (Quorum Technologies Inc.) equipped with a cooled electron-multiplying charge-coupled device (EM-CCD) camera (Hamamatsu) and a 40× oil-immersion objective (NA = 1.3). Cells were maintained at 37 °C with 5% CO₂ throughout imaging.

### Efferocytosis assay

BMDM grown on glass coverslips overnight were exposed to ACs alone or to PA– AC clusters for 30 min followed by sample fixation for microscopy analysis. The Efferocytic Index was calculated as the volume of apoptotic material internalized per cell as described in the Image analysis section.

### Transfection of RAW264.7 cells

RAW264.7 cells were transfected using the Lipofectamine 3000 transfection kit (Thermo Fisher Scientific). For each well of a 12-well plate, 1 µg of plasmid DNA carrying the PM construct and 1 µl of Lipofectamine reagent were diluted in 25 µl RPMI.

### Image Analysis

Images were analyzed using ImageJ (NIH) or Volocity software (PerkinElmer). To quantify the volume of intracellular apoptotic material, the 3D Object Counter plugin for ImageJ was used, as described in[15]. Briefly, this plugin identifies and enumerates fluorescently labeled objects in z-stacks using a user-defined voxel intensity threshold. The plugin generates a binary mask of the objects, where adjacent foreground voxels are considered part of the same particle, and provides a list of particles with their respective volumes (number of voxels). The intracellular versus extracellular localization of particles was determined by visual inspection.

To measure the length of macrophage membrane extensions, the Straight Line tool followed by Analyze ▸ Measure in Fiji (ImageJ) was used. The number of extensions or vesicles per cell was determined by manual counting using the Multi-point tool.

To evaluate the state of internalized apoptotic material, BMDMs were exposed to apoptotic cells at a ratio of one apoptotic cell per three macrophages for varying durations, and the percentage of macrophages containing either a whole internalized apoptotic cell or fragmented apoptotic material was quantified. The diameter of intracellular vesicles containing apoptotic material was measured and compared with the average size of apoptotic cells to distinguish intact cells from fragments.

To quantify the relative surface area occupied by bacteria and apoptotic material within PA–AC clusters, apoptotic cells were labeled with CFSE and bacteria expressed mCherry. Confocal z-stacks were analyzed using Fiji. The surface area of apoptotic material was quantified from the CFSE channel following background subtraction, thresholding, binary image processing, and 3D object analysis. Bacterial surface area was estimated by manually counting bacteria in each cluster and calculating the cumulative surface area based on the average dimensions of a representative sample of bacteria. The ratio between apoptotic material and bacterial surface area was then calculated for each cluster.

When indicated, confocal z-stacks were projected along the z-axis, generating an output image in which each pixel represents the sum of intensities across all z-planes at that x–y position.

### Assessment of PA–AC cluster stability

PA–AC clusters were first allowed to form by incubating *P. aeruginosa* with UV-induced apoptotic cells for 1 h. Following this cluster formation period, stability was assessed by time-lapse microscopy. Individual bacteria associated with apoptotic cells were visually tracked frame by frame over time.

The duration of bacterial tracking depended on the ability to continuously visualize individual bacteria within the field of view, as clusters occasionally moved out of focus or out of the imaging field. Consequently, bacterial association was monitored for a minimum of 15 min and up to a maximum of 30 min.

### Statistical analysis

Statistical analyses of the data were performed using GraphPad Prism 6.0 software (CA, USA). Comparisons among three or more groups were assessed with multiple comparisons test. Comparisons between two groups were assessed using a two-tailed Student’s *t*-test. Differences were considered significant if the p values were < 0.05. For the microscopy studies, error bars correspond to standard errors from three independent experiments.

## Discussion

Our results show that macrophages use distinct modes of engagement when confronted with apoptotic cells alone versus apoptotic cells carrying *P. aeruginosa*. While apoptotic cells can be engulfed as intact structures within well-defined, actin-rich membrane cups, the presence of bacteria alters the geometry of the interaction: macrophages extend actin-rich membrane protrusions that penetrate or skirt gaps in the apoptotic material to reach bacteria embedded within the clusters. Despite the stable attachment of *P. aeruginosa* to the apoptotic surface, these protrusions can extract individual bacteria, which are then internalized into LAMP1-positive, microtubule-associated vesicles. Live-cell imaging confirms that macrophages engage bacteria within clusters directly and are capable of removing them from the apoptotic scaffold.

Quantitative analysis of early vesicles reveals that the majority contain either apoptotic material or bacteria alone, with mixed cargo representing a minority. These findings indicate that, even when apoptotic material and bacteria are presented together as a single, complex target, macrophages can differentiate between these opposing cues and sort their cargo accordingly—a capacity that, to our knowledge, has not been reported before.

Although studies on dual-target phagocytosis are scarce, Naik *et al.* provide an illustrative example using red blood cells opsonized with both IgG and IgM–C3bi. While this model differs from ours—since it features two opsonins coating the same particle rather than distinct targets—they found that simultaneous engagement of the receptors FcγR and CR3 produced additive binding and synergistic uptake, together with a shift toward ruffle-mediated attachment [28]. Jaldin-Fincati *et al*. have also reported that heterogeneous cargo can influence intracellular trafficking. In our study using macrophage-like cells (RAW 264.7 and U937), we examined the uptake and trafficking of aluminium oxyhydroxide adjuvant particles (AlOOH) and the genetically detoxified pertussis toxin (gdPT). When gdPT was adsorbed onto AlOOH, its intracellular fate was altered: rather than escaping the endocytic pathway, the antigen was redirected to degradative endo/lysosomal compartments together with the adjuvant [29].

However, the efferocytosis of bacterially infected apoptotic cells represents the closest precedent to our model, in which macrophages engulf apoptotic cells externally coated with bacteria. In those studies, the pathogens reside within the apoptotic host cell, so that after uptake, the cargo is enclosed within a “double-membrane” structure—the pathogen inside the apoptotic cell, which itself is contained within the macrophage efferosome. These examples show that the spatial configuration of the pathogen can influence vesicular trafficking and maturation: for instance, efferocytosis of *Mycobacterium tuberculosis*–infected apoptotic cells promotes delivery to lysosomes [30] [31]. Dual-target phagocytosis remains largely unexplored, and consequently, so does the impact of target arrangement. In our model, phagocytosis of PA–AC clusters generates three types of vesicles within the same cell—containing bacteria alone, apoptotic material alone, or mixed cargo in which both bacteria and apoptotic material directly contact the vesicle lumen. This configuration provides an opportunity to investigate how macrophages handle these different vesicles, including their trafficking, maturation, and acidification dynamics. It may also allow exploration of how antigen presentation is shaped. This could be particularly intriguing for the mixed vesicles that might present potentially conflicting cues. Apoptotic cargo normally engages LC3-associated phagocytosis, a pathway that promotes rapid maturation while limiting antigen presentation, whereas pathogen-derived signals favor classical phagocytic routes that enhance antigen display [17]. In the mixed vesicles, opposing signals could mislead trafficking, potentially resulting in self-antigen presentation from apoptotic material or reduced presentation of bacterial antigens. Such vesicles thus offer a unique opportunity to investigate how macrophages negotiate conflicting signals and the consequences for cargo processing and immune recognition.

We have determined that macrophages interacting with PA–AC clusters fail to form the actin-rich membrane cups that tightly enclose entire targets, as they do when interacting with apoptotic cells alone. Instead, macrophages adopt membranous extensions that intricately penetrate the apoptotic corpses, thereby creating a conduit to access attached bacteria—and ultimately enabling their internalization. This phenomenon bears some resemblance to the findings of Davidson & Wood, who observed that when macrophages (in the *Drosophila* embryo) encounter apoptotic corpses under high spatial constraint, they employ “filopodial phagocytosis”, in which spike-like filopodia extend to draw in debris. In contrast, under low spatial constraints they preferentially use “lamellipodial phagocytosis”, in which broad sheet-like lamellipods envelop the target [32].

Our results also show that when macrophages encounter bacteria-coated apoptotic cells, the apoptotic cells are fragmented before complete engulfment, reflecting a piecemeal uptake mode. Interestingly, studies of phagocytosis involving geometrically complex targets indicate that phagocytes adjust their engulfment strategy in response to local curvature cues, such as shifting from complete ingestion of the target to partial uptake [33].

Our observation that membrane protrusions engaging and internalizing *P. aeruginosa* align with microtubules, and that Lamp1-positive vesicles containing bacterial cargo are distributed along these microtubule-rich extensions, resonates with recent findings in microglia describing two distinct modes of phagocytosis [34]. In that study, the authors reported a predominant *branch-mediated* mode, in which phagosomes form at the tips of long, microtubule-containing branches and are subsequently transported toward the cell body, and a secondary *non-branch-mediated* mode occurring directly at the soma.

Disruption of microtubules induced a shift from the branch-mediated to the non-branch-mediated mode, underscoring the dependence of distal phagocytosis on microtubule integrity. Moreover, centrosome positioning toward the active branch was shown to facilitate phagosome formation and transport. Although our system differs substantially, both studies point to the possibility that microtubule organization supports distal forms of phagocytosis, enabling uptake events to occur at sites distant form the cell body. Consistent with this concept, our finding that microtubule disruption impairs PA–AC cluster internalization supports a role for microtubules in regulating protrusion-based uptake. In the future, it will be interesting to investigate how microtubules contribute to phagocytosis of PA–AC clusters.

Altogether, this work uncovers a previously unrecognized facet of macrophage phagocytic plasticity and reveals that these cells can differentially engage and sort the components of complex dual targets. The molecular mechanisms underlying this selective response remain to be elucidated. Future studies will be required to determine how macrophages integrate signals derived from distinct cargo components and coordinate the cellular processes that enable differential cargo handling. Thus, this work lays the groundwork for further investigations into how phagocytes decode and respond to complex targets encountered in diverse physiological and pathological contexts.

## Acknowledgments

We thank Dr. Terebiznik and Dr. C. Gimenez for their guidance and support during C.D.’s time in their laboratory at the University of Toronto Scarborough, ON, Canada.. C.D. is a research fellow at CONICET. S.L.M and N.N.R are undergraduated students at UNSAM. A.K. is a research career member of CONICET.

## Supporting information

**S1 Video. Live-cell time-lapse of a BMDM exposed to *P. aeruginosa*.**

**S2 Video. Stability of PA–AC clusters**

Time-lapse of preformed PA–AC clusters, with each frame presented as a maximum-intensity Z projection. Bacteria remained enclosed within the apoptotic cells throughout the recording. Cyan: Cell Trace-labeled apoptotic cell; red: mCherry-*P. aeruginosa*. Scale bar: 2 μm

**S3 Video. Z-stack visualization displaying sequential XY corresponding to the data shown in Fig 4A**.

**S4 Video . Z-stack visualization displaying sequential XY corresponding to the data shown in Fig 4B**.

**S5 Video. Z-stack visualization displaying sequential XY corresponding to the data shown in Fig 4C**.

**S6 Video. Z-stack visualization displaying sequential XY corresponding to the data shown in Fig 5A (lower panel).**

**S7 Video. Z-stack visualization displaying sequential XY corresponding to the data shown in Fig 5A (upper panel).**

**S8 Video. Macrophage membrane remodeling during bacterial uptake from a PA–AC cluster**

Live-cell time-lapse confocal video of RAW 264.7 macrophages expressing a plasma membrane marker, incubated at a 1:5 ratio with PA–AC clusters. Gray: PM-GFP, plasma membrane marker; green: Cell Trace-labeled apoptotic cells; red: mCherry-*P. aeruginosa*.

**S9 Video. Macrophage actin remodeling during bacterial uptake from a PA–AC cluster**

Live-cell time-lapse confocal video of RAW 264.7 macrophages expressing LifeAct-RFP, incubated at a 1:5 ratio with PA–AC clusters. Red: LifeAct-RFP, F-actin marker; gray: Cell mask-labeled apoptotic cells; green: mCherry-*P. aeruginosa*.

## References

1. Cockram TOJ, Dundee JM, Popescu AS, Brown GC. The Phagocytic Code Regulating Phagocytosis of Mammalian Cells. Frontiers in Immunology. 2021. doi:10.3389/fimmu.2021.629979

2. Flannagan RS, Jaumouillé V, Grinstein S. The cell biology of phagocytosis. Annual Review of Pathology: Mechanisms of Disease. 2012. doi:10.1146/annurev-pathol-011811-132445

3. Franc NC, White K, Ezekowitz RAB. Phagocytosis and development: Back to the future. Curr Opin Immunol. 1999;11. doi:10.1016/S0952-7915(99)80009-0

4. Pauwels AM, Trost M, Beyaert R, Hoffmann E. Patterns, Receptors, and Signals: Regulation of Phagosome Maturation. Trends in Immunology. 2017. doi:10.1016/j.it.2017.03.006

5. Birge RB, Boeltz S, Kumar S, Carlson J, Wanderley J, Calianese D, et al. Phosphatidylserine is a global immunosuppressive signal in efferocytosis, infectious disease, and cancer. Cell Death and Differentiation. 2016. doi:10.1038/cdd.2016.11

6. Boada-Romero E, Martinez J, Heckmann BL, Green DR. The clearance of dead cells by efferocytosis. Nature Reviews Molecular Cell Biology. 2020. doi:10.1038/s41580-020-0232-1

7. Rieber N, Hector A, Carevic M, Hartl D. Current concepts of immune dysregulation in cystic fibrosis. Int J Biochem Cell Biol. 2014;52: 108–112. doi:10.1016/j.biocel.2014.01.017

8. Soleti R, Porro C, Martinez MC. Apoptotic process in cystic fibrosis cells. Apoptosis. 2013/06/25. 18: 1029–1038. doi:10.1007/s10495-013-0874-y

9. Schwarzer C, Fischer H, Machen TE. Chemotaxis and Binding of Pseudomonas aeruginosa to Scratch-Wounded Human Cystic Fibrosis Airway Epithelial Cells. PLoS One. 2016;11: e0150109. doi:10.1371/journal.pone.0150109

10. Martin CJ, Peters KN, Behar SM. Macrophages Clean Up: Efferocytosis and Microbial Control The Many Jobs of the Macrophage. Curr Opin Microbiol. 2014;0.

11. Mohammad-Rafiei F, Moadab F, Mahmoudi A, Navashenaq JG, Gheibihayat SM. Efferocytosis: a double-edged sword in microbial immunity. Archives of Microbiology. 2023. doi:10.1007/s00203-023-03704-8

12. Rosenblatt J, Raff MC, Cramer LP. An epithelial cell destined for apoptosis signals its neighbors to extrude it by an actin- and myosin-dependent mechanism. Curr Biol. 2001;11: 1847–1857.

13. Capasso D, Pepe M V, Rossello J, Lepanto P, Arias P, Salzman V, et al. Elimination of Pseudomonas aeruginosa through Efferocytosis upon Binding to Apoptotic Cells. PLoS Pathog. 2016;12: e1006068. doi:10.1371/journal.ppat.1006068

14. Pepe MV, Dea C, Genskowsky C, Capasso D, Roset MS, Jäger AV, et al. Reversible adhesion by type IV pili leads to formation of permanent localized clusters. iScience. 2022;25. doi:10.1016/J.ISCI.2022.105532

15. Jäger AV, Arias P, Tribulatti MV, Brocco MA, Pepe MV, Kierbel A. The inflammatory response induced by Pseudomonas aeruginosa in macrophages enhances apoptotic cell removal. Sci Rep. 2021;11. doi:10.1038/s41598-021-81557-1

16. Nakaya M, Kitano M, Matsuda M, Nagata S. Spatiotemporal activation of Rac1 for engulfment of apoptotic cells. Proc Natl Acad Sci U S A. 2008;105. doi:10.1073/pnas.0803677105

17. Yin C, Heit B. Cellular Responses to the Efferocytosis of Apoptotic Cells. Frontiers in Immunology. 2021. doi:10.3389/fimmu.2021.631714

18. Vorselen D. Dynamics of phagocytosis mediated by phosphatidylserine. Biochemical Society Transactions. 2022. doi:10.1042/BST20211254

19. Hoijman E, Häkkinen HM, Tolosa-Ramon Q, Jiménez-Delgado S, Wyatt C, Miret-Cuesta M, et al. Cooperative epithelial phagocytosis enables error correction in the early embryo. Nature. 2021;590. doi:10.1038/s41586-021-03200-3

20. Raymond MH, Davidson AJ, Shen Y, Tudor DR, Lucas CD, Morioka S, et al. Live cell tracking of macrophage efferocytosis during Drosophila embryo development in vivo. Science (1979). 2022;375. doi:10.1126/science.abl4430

21. Yin C, Argintaru D, Heit B. Rab17 mediates intermixing of phagocytosed apoptotic cells with recycling endosomes. Small GTPases. 2019;10. doi:10.1080/21541248.2017.1308852

22. Samie M, Wang X, Zhang X, Goschka A, Li X, Cheng X, et al. A TRP channel in the lysosome regulates large particle phagocytosis via focal exocytosis. Dev Cell. 2013;26. doi:10.1016/j.devcel.2013.08.003

23. Searle S, Bright NA, Roach TIA, Atkinson PGP, Barton CH, Meloen RH, et al. Localisation of Nramp1 in macrophages: Modulation with activation and infection. J Cell Sci. 1998;111. doi:10.1242/jcs.19.111.2855

24. Falcón-Pérez JM, Nazarian R, Sabatti C, Dell’Angelica EC. Distribution and dynamics of Lamp1-containing endocytic organelles in fibroblasts deficient in BLOC-3. J Cell Sci. 2005;118. doi:10.1242/jcs.02633

25. Teruel MN, Blanpied TA, Shen K, Augustine GJ, Meyer T. A versatile microporation technique for the transfection of cultured CNS neurons. J Neurosci Methods. 1999;93. doi:10.1016/S0165-0270(99)00112-0

26. Marim FM, Silveira TN, Lima DS, Zamboni DS. A Method for Generation of Bone Marrow-Derived Macrophages from Cryopreserved Mouse Bone Marrow Cells. PLoS One. 2010;5: e15263. doi:10.1371/JOURNAL.PONE.0015263

27. Lagendijk EL, Validov S, Lamers GE, de Weert S, Bloemberg G V. Genetic tools for tagging Gram-negative bacteria with mCherry for visualization in vitro and in natural habitats, biofilm and pathogenicity studies. FEMS Microbiol Lett. 2010;305: 81–90. doi:10.1111/j.1574-6968.2010.01916.x

28. Naik U, Nguyen QPH, Harrison RE. Binding and uptake of single and dual-opsonized targets by macrophages. J Cell Biochem. 2020;121. doi:10.1002/jcb.29043

29. Jaldin-Fincati J, Moussaoui S, Gimenez MC, Ho CY, Lancaster CE, Botelho R, et al. Aluminum hydroxide adjuvant diverts the uptake and trafficking of genetically detoxified pertussis toxin to lysosomes in macrophages. Mol Microbiol. 2022;117. doi:10.1111/mmi.14900

30. Martin CJ, Booty MG, Rosebrock TR, Nunes-Alves C, Desjardins DM, Keren I, et al. Efferocytosis is an innate antibacterial mechanism. Cell Host Microbe. 2012;12: 289–300. doi:10.1016/j.chom.2012.06.010

31. Karaji N, Sattentau QJ. Efferocytosis of pathogen-infected cells. Frontiers in Immunology. 2017. doi:10.3389/fimmu.2017.01863

32. Davidson AJ, Wood W. Macrophages Use Distinct Actin Regulators to Switch Engulfment Strategies and Ensure Phagocytic Plasticity In Vivo. Cell Rep. 2020;31. doi:10.1016/j.celrep.2020.107692

33. Clarke M, Engel U, Giorgione J, Müller-Taubenberger A, Prassler J, Veltman D, et al. Curvature recognition and force generation in phagocytosis. BMC Biol. 2010;8. doi:10.1186/1741-7007-8-154

34. Möller K, Brambach M, Villani A, Gallo E, Gilmour D, Peri F. A role for the centrosome in regulating the rate of neuronal efferocytosis by microglia in vivo. Elife. 2022;11. doi:10.7554/eLife.82094

